# Genetically-identified cell types in avian pallium mirror core principles of excitatory and inhibitory neurons in mammalian cortex

**DOI:** 10.1101/2020.11.11.374553

**Authors:** Jeremy A. Spool, Matheus Macedo-Lima, Garrett Scarpa, Yuichi Morohashi, Yoko Yazaki-Sugiyama, Luke Remage-Healey

## Abstract

In vertebrates, advanced cognitive abilities are associated with a highly developed telencephalic pallium. In mammals, the six-layered neocortex of the pallium is composed of excitatory neurons and inhibitory interneurons, organized across layers into microcircuits. These organizational principles are proposed to support efficient, high-level information processing. Comparative perspectives across vertebrates provide a lens to understand what common features of pallium are important for complex cognition. For non-mammalian vertebrates that exhibit complex cognitive abilities, such as birds, the physiology of identified pallial cell types and their circuit organization are largely unresolved. Using viral tools to target excitatory vs. inhibitory neurons in the zebra finch auditory association pallium, we systematically tested predictions derived from mammalian neocortex. We identify two segregated neuronal populations that exhibit profound physiological and computational similarities with mammalian excitatory and inhibitory neocortical cells. Specifically, despite dissimilarities in gross architecture, avian association pallium exhibits neocortex-typical coding principles, and inhibitory-dependent cortical synchrony, gamma oscillations, and local suppression. Our findings suggest parallel evolution of physiological and network roles for pallial cell types in amniotes with substantially divergent pallial organization.

## Main Text

The vertebrate pallium is considered essential for advanced cognitive abilities. Unlike other sectors of the vertebrate brain, the cytoarchitecture and elaboration of pallial domains vary immensely across vertebrate classes, and even within closely related taxa^1^. In mammals, for example, dorsal pallium develops into a six-layered structure called neocortex, the elaboration of which is associated with advanced cognitive abilities^2,3^. In cartilaginous fishes, some taxa have a layered pallium, while in other species the organization of pallium is not at all obvious, and difficult to separate from subpallial structures^1^. Large swaths of bird pallium are unlayered^4,5^ (Fig. 1A; though see ^6–9^ for evidence of lamination in auditory pallium and hyperpallium), yet several species of birds, including parrots and corvids, exhibit cognitive abilities on par or exceeding capabilities of our closest primate relatives^10^. Deep characterization of the molecular properties and developmental origins of pallial cells has advanced our understanding of pallial evolution (e.g.^11–18^). However, a key piece missing is an examination of the physiology of pallial cell types in non-mammalian species. In this study, we begin to address this gap by interrogating the physiological and network phenotypes of two cell types in avian auditory association pallium.

**Fig. 1.**
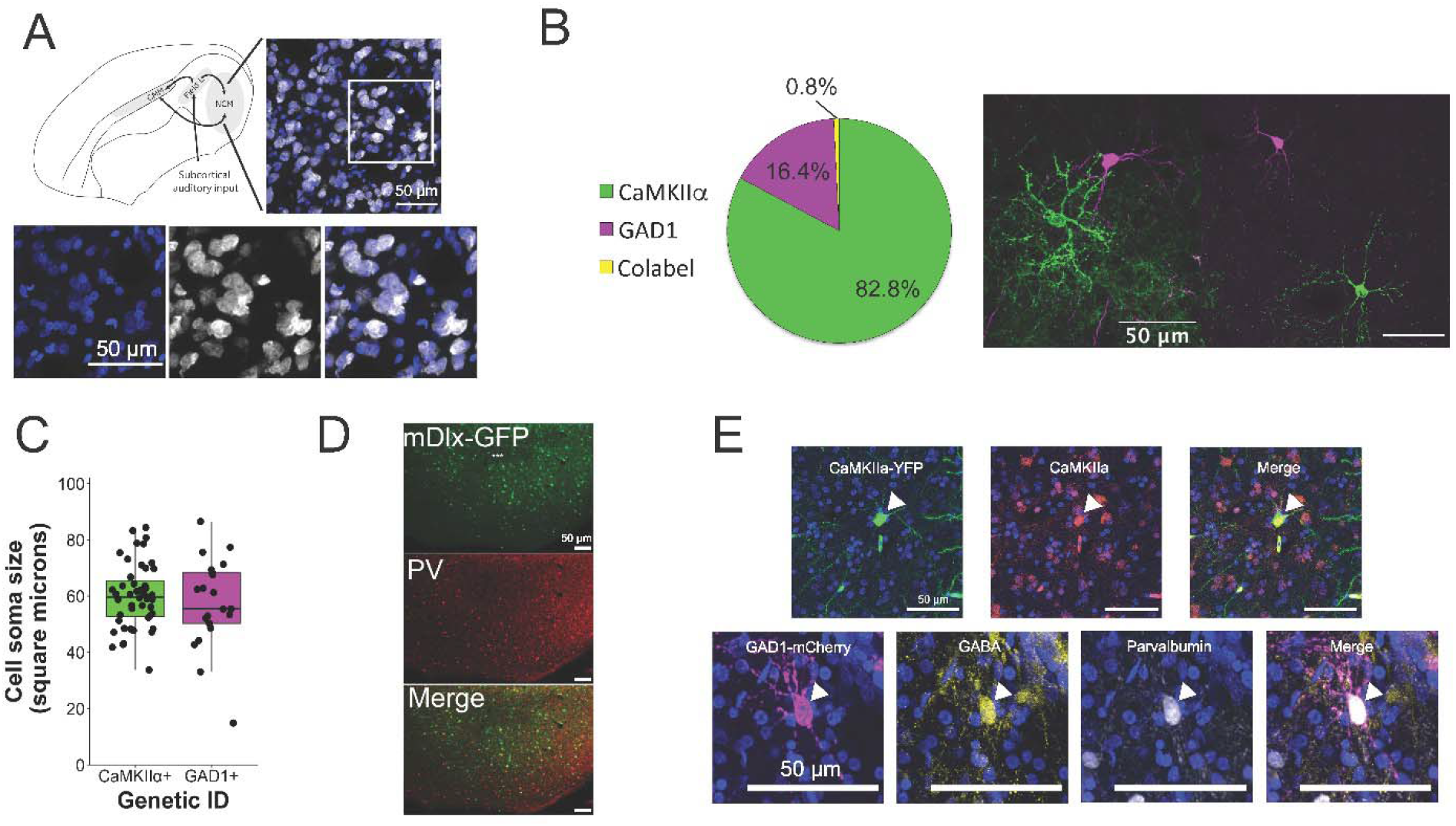
Viruses targeting CaMKIIα and GAD1 promoters segregate cell types in avian auditory association pallium. (**A**) Sagittal schematic of avian pallium (top left) showing primary auditory cortex (Field L), and auditory association pallium (caudomedial mesopallium, CMM; caudomedial nidopallium, NCM). In the schematic, left is rostral, right is caudal, top is dorsal, and bottom is ventral. 20x confocal image (top right) shows non-laminar clustered cytoarchitecture of NCM. Bottom: magnified from white box in top right (blue=DAPI; white=NeuN). (**B**) Non-overlapping cell types identified by viral expression of fluorophores (left), and 60x images of transduced CaMKIIα (green) and GAD1 (magenta) cells. (**C**) Cell soma area at largest cross-sectional diameter. (**D**) Typical widefield view of viral injection site in avian NCM. Shown is mDlx-GFP expression (top), parvalbumin immunolabeling (PV; middle), and overlaid images (bottom) showing specificity (>66% viral cells co-labeled) and efficiency (>88% PV cells co-labeled) of transduction. (**E**) 60x images of viral and antibody co-expression. Top: CaMKIIα viral expression (green) colocalizing with CaMKIIα immunolabeling (orange). Bottom: GAD1 viral expression (magenta) co-localizing with GABA (gold) and PV (white). White arrowheads = co-localization. All scale bars = 50 microns.

While there remains a lack of consensus over how the majority of the avian pallium is related to mammalian pallium, there are unquestionable similarities in pallial circuit organization and function between the two taxa. The adult function and connectivity of the avian pallium appears highly similar to mammalian neocortex, including thalamic input to primary pallial structures, high interconnectivity of pallial regions, and output to subpallial brain structures^19–23^. Brain regions in the avian pallium perform functions of similar cognitive complexity to functions ascribed to neocortex, including but not limited to encoding memory of complex, ecologically-relevant sensory stimuli, vocal learning, encoding numerical information, cross modal associative learning, and decision-making based on abstract rules^24–33^. Brain molecular expression analyses have identified striking similarities between birds and mammals at the level of pallial cell types^12,17,34^. The physiology of pallial cell types in birds is unresolved. Currently, the identification of cell types strongly resembling mammalian cortical cell-types in avian pallium is putative. For example, extracellular recordings in avian pallium consistently identify broad-spiking neurons with sparse firing rates that have selective receptive fields for ecologically-relevant stimuli, and narrow-spiking neurons with higher firing rates that are less selective among stimuli^7,28,35–42^. Such similar circuit organization and function could reflect homologous structures and cell types in birds and mammals, convergent evolution of cell types originating from nearby pallial compartments, or parallel evolution of circuit features from basal cell types that existed in the last common ancestor of amniotes.

A definitive mapping of the physiology of pallial cells onto their molecular identity in birds is critical for understanding the extent to which synaptic and computational properties track with molecular phenotype compared to mammalian neocortical circuits. To date, physiological dissection of non-mammalian pallium has been limited by a lack of tools to precisely identify and control specific cell types. Here, we deployed viral tools in birds to dissect circuits in a secondary auditory associative region of pallium, the caudomedial nidopallium (NCM; Fig. 1A). NCM is highly interconnected, is involved in auditory memory and individual recognition, and the auditory physiology of neurons in this region is well-described^28,35–37,39,40^. We identified two classes of neurons with shared physiological features to neocortical excitatory neurons vs.

inhibitory interneurons in mammals. Specifically, despite dissimilarities in anatomical organization, this study reveals shared intrinsic physiological properties, auditory coding roles, and network organization in which inhibition drives local suppression and synchronizes large-scale gamma oscillations. The extent of these similarities in birds and mammals may provide insight into what pallial circuit features support complex cognition in amniotes.

## Results

We established methods for viral optogenetics in the NCM of zebra finches (*Taeniopygia guttata*) to test predictions derived from mammalian neocortical excitatory neurons vs. inhibitory interneurons. In mammalian cortex, markers like calmodulin-dependent kinase (CaMKIIα) identify excitatory neurons, while the GABA-producing enzyme glutamate decarboxylase 1 (GAD1) and the homeobox-containing mDlx gene identify inhibitory neurons^43–5^. We first injected zebra finch NCM with adeno-associated viruses driving opsin proteins under the control of the CaMKIIα vs. GAD1 promoters, and observed clear segregation of transfected cells (Fig. 1B). The ratio of CaMKIIα vs. GAD1 abundance was 83:16, echoing the distribution of excitatory and inhibitory cell types in mammalian neocortex (GABAergic neurons typically comprise ~10-20% of cortical neurons, e.g.^46^). Qualitatively, we observed that CaMKIIα cells were morphologically variable with respect to dendritic spine density and thickness of dendritic branches, while GAD1 cells were more often aspiny, with thinner processes (fig. S1). CaMKIIα cells and GAD1 cells did not differ in soma size (Welch’s t_29.4_ = 0.66, *P* = 0.51; Fig. 1C). Conventional antibody staining confirmed selective transduction of cell-type targets (Fig. 1D,E). Thus, promoter-driven molecular cell identity segregates cell types in avian association pallium as it does in mammalian neocortex.

We next examined the physiology of identified cell types in zebra finch NCM. Mammalian excitatory neocortical neurons typically have phasic, accommodating responses to depolarizing current injections and broad spike widths, whereas neocortical interneurons have tonic responses and narrow spike widths^47,48^. In NCM whole-cell slice recordings (Fig. 2A), depolarizing current steps elicited phasic firing profiles from CaMKIIα cells vs. tonic firing profiles from GAD1 cells (Fig. 2B,C). Similar to phasic excitatory neurons in mammalian barrel and other cortices^49–51^, CaMKIIα cells responded with maximum 1-2 spikes (ISIs 48.8 +/- 26.6 ms; range 20-114 ms), in contrast to the tonically-responsive, non-accomodating profile of GAD1 cells (ISIs 48.2 +/- 11.4 ms; range 35-70 ms) that exhibit very low adaptation ratios^52^: 1.105 +/- 0.02. CaMKIIα cells also had broader action potential widths compared to GAD1 cells (Mann-Whitney test: W = 162, *P* = 0.005; Fig. 2D), but did not differ in other passive membrane properties (input resistance: Welch’s t_20.8_ = −1.4728, *P* = 0.1558; rheobase: t_23.8_ = 1.3593, *P* = 0.1868; Fig. 2E,F). Next, we conducted *in vivo* optrode recordings to isolate properties of photoidentified CaMKIIα vs. GAD1 single units (Fig. 2G). Spike widths were broader in CaMKIIα units than GAD1 units (action potential width (peak-to-peak): W = 318, *P* = 2.7e-05; Fig. 2H; width of action potential at quarter height (i.e., spike quarter-width): W = 342, *P* = 9.9e-07; Fig. 2I; no difference in non-light evoked units; fig. S2). Light-evoked spike latencies were longer for CaMKIIα compared to GAD1 single units (W = 273.5, *P* = 0.004; Fig. 2J), consistent with their slower spike onset kinetics (fig. S3) and greater degree of network suppression. The sharp physiological distinctions between CaMKIIα and GAD1 neurons in NCM therefore mirror those of mammalian neocortical excitatory neurons and inhibitory interneurons, respectively.

**Fig. 2.**
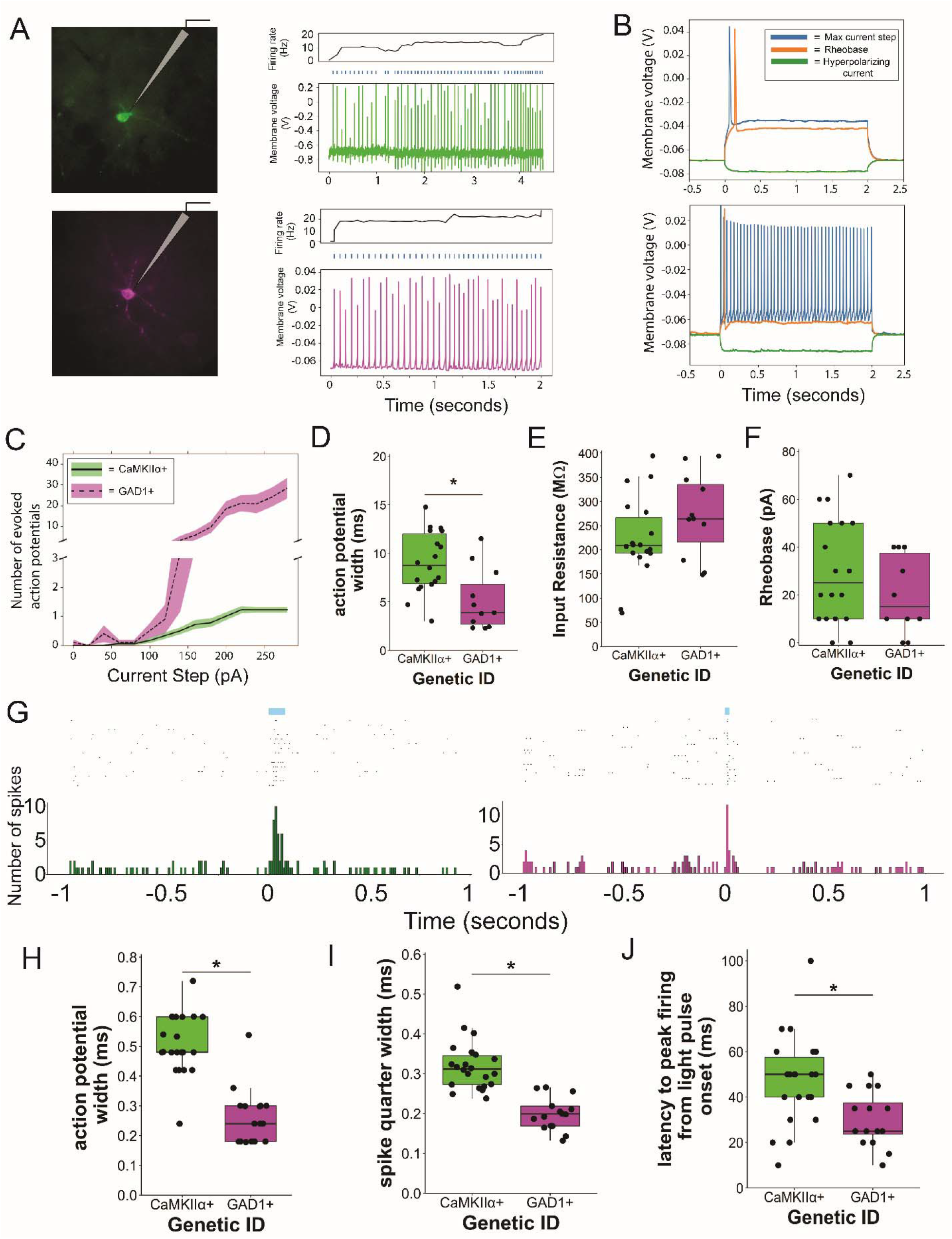
CaMKIIα and GAD1 single units in NCM have distinct physiological properties. (**A**) Transduced cells in whole-cell current clamp configuration (left) and reliable photopotentials to 25 ms blue light pulses (top row = CaMKIIα cell; bottom row = GAD1 cell). (**B**) CaMKIIα cells exhibit phasic responses (top) while GAD1 cells exhibit tonic responses (bottom) to current steps. (**C**) Mean + SEM action potentials for CaMKIIα cells (green; n = 18) and GAD1 cells (magenta; n = 11) in response to current steps. (**D**) Spike width, (**E**) Input resistance, and (**F**) Rheobase of CaMKIIα cells vs. GAD1 cells in whole-cell recordings. (**G**) Raster plots and histograms of exemplar *in vivo* transduced single units in electrophysiological response to blue light pulses. (**H**) Action potential widths, (**I**) spike quarter-widths, and (**J**) latency to light-evoked response peak in optically-identified single units *in vivo. *P* < 0.05 for Mann-Whitney U tests.

Auditory coding roles are distinct between cell types in mammalian neocortex, where auditory stimuli elicit higher spiking activity from interneurons and quicker latencies compared to sparse-firing excitatory cells^53–55^. We therefore tested whether *in vivo* responses of CaMKIIα vs. GAD1 neurons segregated accordingly. Conspecific song drove GAD1 at higher rates compared to CaMKIIα units (Fig. 3A,B; W = 106, *P* = 0.040). GAD1 single units had a quicker response latency compared to CaMKIIα units (Fig. 3C; W = 207, *P* = 0.002). In rat auditory cortex, inhibitory interneurons typically respond promiscuously to sensory stimuli, whereas excitatory neurons in association layers have more selective representations^55^. Likewise in associative auditory avian pallium, putative excitatory neurons are more stimulus-selective compared to putative interneurons^37^. Single-unit spike trains fed to a custom pattern classifier produced higher decoding accuracy values for GAD1 than for CaMKIIα units, indicating that GAD1 units carried information about a variety of auditory stimuli (W = 84, *P* = 0.006825; Fig. 3D,E; fig. S4). By contrast, when examining stimulus selectivity (see Materials and Methods), CaMKIIα single units tended to be more selective for a subset of conspecific song stimuli (W = 241, *P* = 0.057; Fig. 3D,F). These distinctions in auditory coding roles for CaMKIIα and GAD1 neurons precisely match the established sensory encoding roles of mammalian neocortical excitatory neurons vs. inhibitory interneurons.

**Fig. 3.**
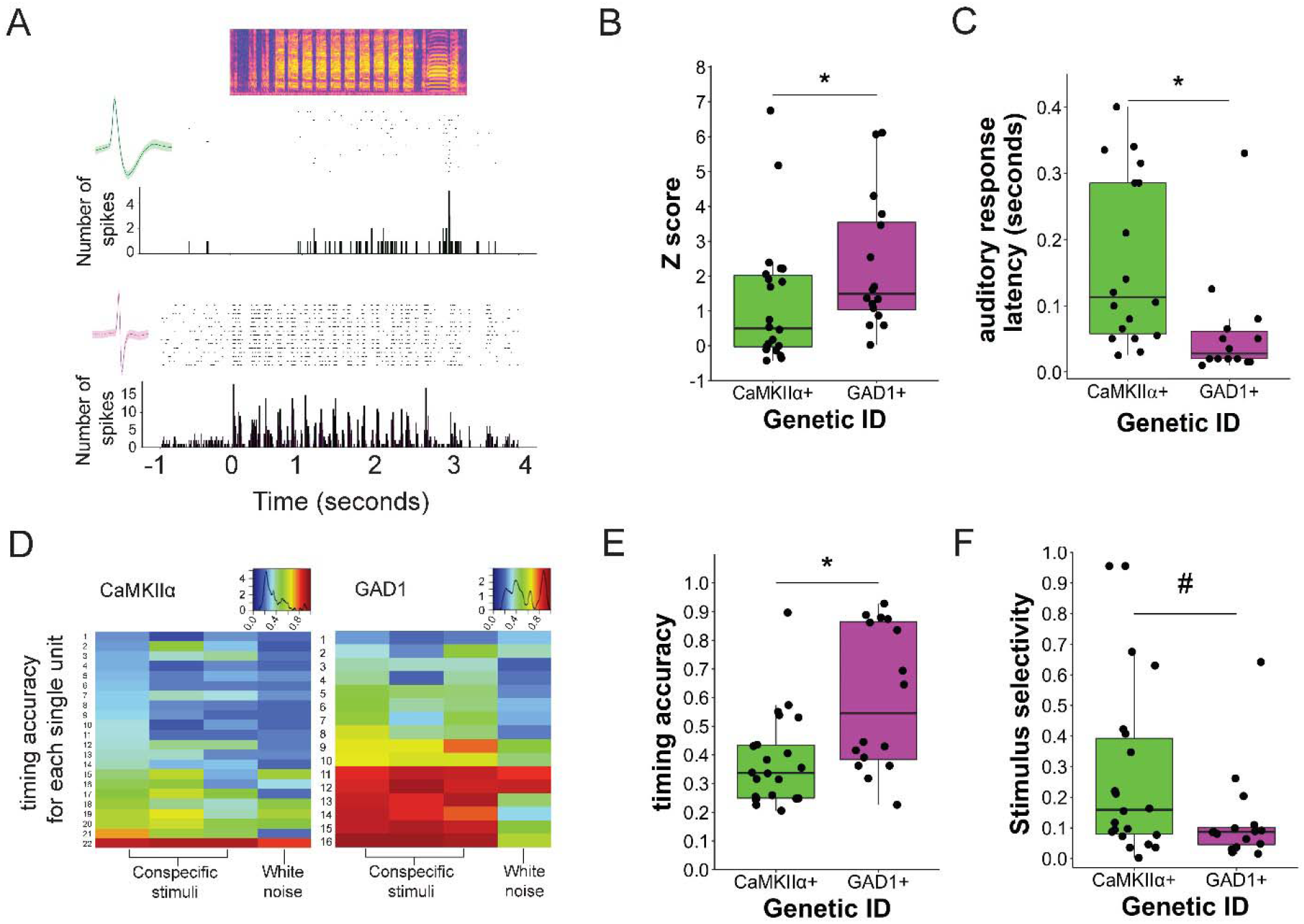
Auditory-response properties of CaMKIIα and GAD1 neurons. (**A**) Spectograms and rasterplots of a CaMKIIα single unit (top) and a GAD1 single unit (bottom) in NCM responding to conspecific song. (**B**) Song-evoked Z scores of optically-identified single units. (**C**) Latency in seconds for optically-identified single units to respond to white noise stimuli. (**D**) Heat map of single unit timing accuracy measures across auditory stimuli. Values closer to 1 represent higher timing accuracy. Histogram insets show density distribution of accuracy metric across stimuli for CaMKIIα (left) and GAD1 single units (right). (**E**) Pattern classifier timing accuracy averaged across auditory stimuli for optically-identified single units displayed in D. (**F**) Measure of transduced single unit selectivity for subsets of conspecific stimuli (see Materials and Methods). *\*P* < 0.05 and *#P* = 0.057 for Mann-Whitney U tests.

One well-described feature of mammalian neocortical microcircuits is feedforward suppression, in which incoming excitation drives inhibitory interneurons followed by excitatory neurons in tandem, yielding temporally-precise excitation quenched rapidly by inhibition^53,56,57^.

Consistent with this prediction, GAD1 neurons had faster auditory onset latency than CaMKIIα neurons (Fig. 3C). Furthermore, anesthetized optrode recordings showed that GAD1-ChR2 optical stimulation drove short-latency suppression in ~25% of non-transduced single units, compared to 0% for CaMKIIα-ChR2 experiments (fig. S5). To test this further in awake animals, we directed a 32-channel opto-microdrive at the NCM of mDlx-ChR2-transduced birds. At sites with photo-identified inhibitory neurons, a separate population of cells were photo-suppressed and quickly rebounded at light-pulse offset (Fig. 4A). We also identified units that rebounded at light-pulse offset using a GAD1-ChR2 construct in an awake animal (fig. S6A). mDlx and GAD1 interneurons therefore suppress local synaptic targets in avian NCM, in a manner similar to mammalian neocortex.

**Fig. 4.**
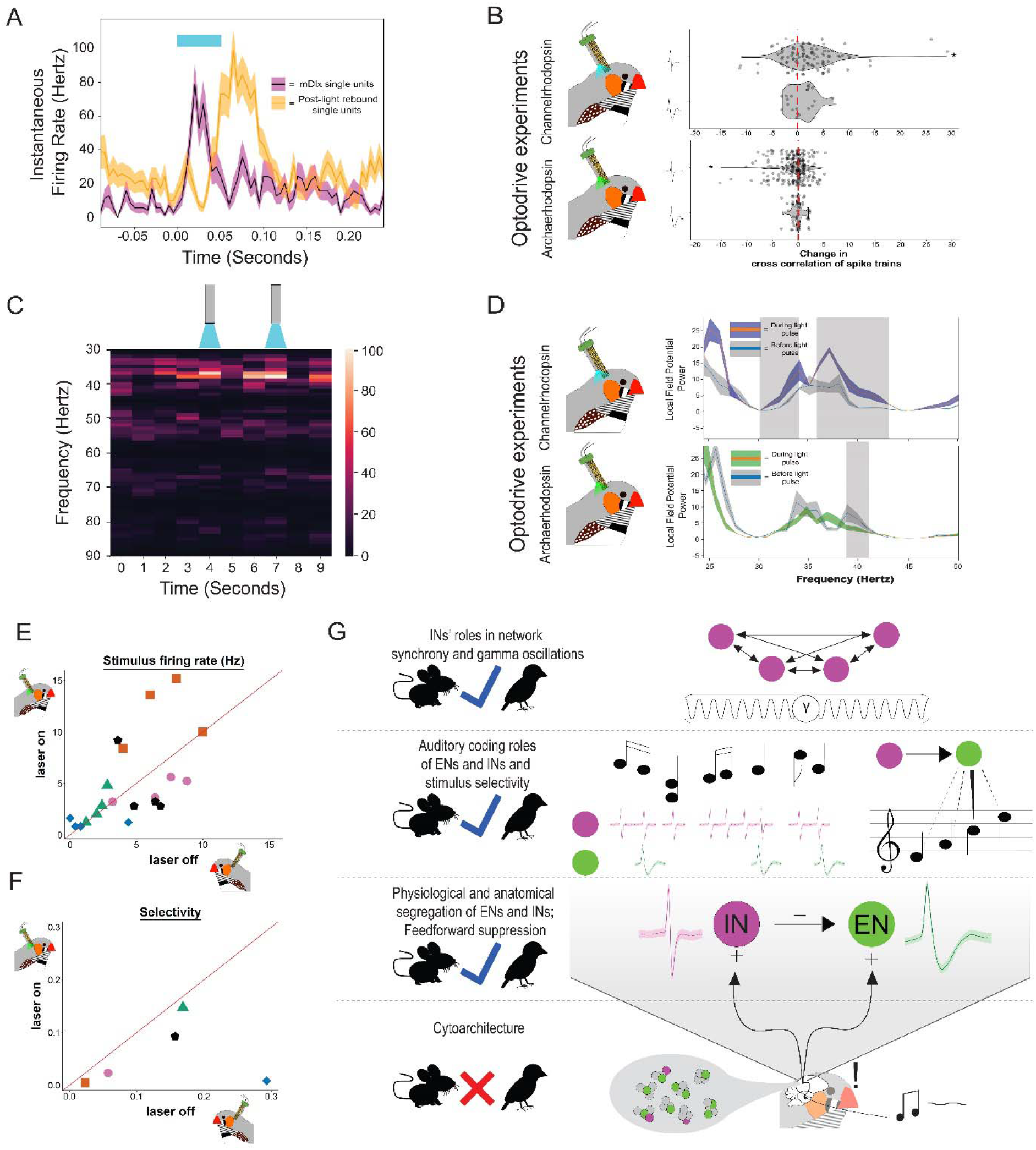
Inhibitory neurons in NCM drive suppression, synchrony, gamma oscillations, and functional auditory response properties. (**A**) Instantaneous firing rates of light-evoked single units (magenta; n = 5) and light-suppressed single units (orange; n = 5) from optical stimulation (50 ms blue pulse) of mDlx-ChR2-transduced cells drawn from n = 36 total units isolated in NCM *in vivo*. (**B**) Violin plots show change in cross-correlation of waveforms when NCM was stimulated with blue light in mDlx-ChR2 experiments (top row) and when stimulated with green light for GAD1-archaerhodopsin experiments (bottom). Red line denotes zero change. (**C**) Heat map (% max LFP power) showing LFP change over time in an mDlX-ChR2 optodrive experiment. Cartoon lasers above represent bins with blue light pulses. (**D**) LFP power spectra before and during stimulation of NCM with blue light for mDlx-ChR2 experiments (top) and with green light for GAD1-archaerhodopsin experiments (bottom). Gray shading represents gamma frequency range for which LFP power is significantly different from baseline according to predictions from mammalian cortex (*P* < 0.05); $ is range for which LFP power is significantly different from baseline against predicted direction (*P* < 0.05). (**E**) Single units with broad waveforms are plotted with respect to their stimulus firing rates in hertz to various conspecific songs when GAD1-archaerhodopsin was stimulated with green laser (y-axis) compared to when there was no laser (x-axis). Each point represents a single unit’s response to one conspecific song, and single units are grouped by a unique shape and color. (**F**) The same single units as E are plotted with respect to their selectivity across all conspecific stimuli when green laser was on (y-axis) compared to when green laser was off. For E & F, points deviating from the red line denote a non-zero difference between laser off and laser on conditions. (**G**) Summary schematic showing features of avian excitatory principal cell-like (EN) and inhibitory interneuron-like (IN) neurons in NCM in reference to predictions from mammalian cortex. Arrows imply effects of neuron stimulation and do not imply nature of synaptic connectivity.

Inhibitory interneurons in mammalian neocortex also coordinate broad-scale spike synchrony and oscillations in the gamma frequency band^58,59^. With mDlx-ChR2, optical stimulation of NCM increased synchrony of single units with narrow waveforms (narrow single units, W = 4184, *P* = 2e-05; broad single units, W = 146, *P* = 0.124; Fig. 4B). Optical stimulation of mDlx cells in NCM also increased the amplitude of the gamma frequency band at 30-34 and 36-43 Hz (All *P* < 1e-05; Fig. 4C,D). Optodrive recordings with a GAD1-ChR2 construct showed similar results (fig. S6B,C). Conversely, with a GAD1-archaerhodopsin construct that hyperpolarized avian cells *in vitro* (fig. S7), we observed that optical suppression of GAD1-archaerhodopsin cells decreased synchrony of narrow but not broad cells (narrow: W = 5288.5, *P* = 6e-04; broad: W = 206, *P* = 0.95; Fig. 4B) and attenuated the gamma band specifically at 39-41 Hz (All *P* < 0.001; Fig. 4C,D). Rapid shifts in NCM inhibitory network activation therefore drove tuning of cortical network synchrony and gamma oscillations.

Finally, a balance of excitation and inhibition is thought to be critical for neocortical function^60^. In rodent auditory neocortex, disrupting inhibition alters excitatory neuron auditory responses, including firing rates, and stimulus selectivity^61^. To test whether this was the case in zebra finch auditory pallium, we examined optodrive recordings using GAD1-archaerhodopsin in NCM. Specifically, we recorded the responses of broad waveforms to conspecific songs with and without green laser stimulation. Firing rates during stimulus presentations in single units with broad waveforms were not different as a group between laser off and laser on conditions (Fig. 4E; Wilcoxon signed rank test, V = 55; p = 0.80; Chi-square test for increased/decreased firing, X_1_ = 0.8, p = 0.371). However, all single units with broad waveforms had decreased stimulus selectivity during laser on vs. laser off conditions (Fig. 4F; Wilcoxon signed rank test, V = 15; p = 0.063; Chi-square test for increased/decreased selectivity, X_1_ = 5, p = 0.025). This suggests that disruption of inhibition in NCM has functional consequences for coding of ethologically-relevant auditory stimuli.

## Discussion

Our findings demonstrate compelling computational and physiological similarities between corresponding excitatory and inhibitory neuronal cell types of avian association pallium and mammalian neocortex (Fig. 4G). Viruses designed to target excitatory neurons and inhibitory interneurons in mammalian neocortex segregate similar populations in avian association pallium with predicted intrinsic physiology, auditory coding roles, and network organization that drives local suppression and synchronizes large-scale gamma oscillations, all despite a non-laminar organization.

The mapping of physiological roles to molecular identity of excitatory and inhibitory neurons in pallium is not a generalized feature of the vertebrate or even the mammalian brain. For example, in contrast to pallium, GABAergic medium spiny neurons in the striatum express high levels of CaMKIIα, and are the major source of efferent projections from the region^62,63^, and both projection and interneurons in the striatum release GABA^63,64^. In the lateral hypothalamus, parvalbumin-positive, glutamatergic neurons are fast-spiking and send long-range projections in the brain^65^ Relative proportions of glutamatergic vs. GABAergic cell populations also vary between brain regions, including but not limited to the amygdala, bed nucleus of the stria terminalis, and ventral tegmental area, where glutamatergic neurons are the minority^66–68^.

Neurons in some brain regions can also co-release GABA and glutamate^69,70^. Therefore, there are many ways the brain builds functional circuits using all combinations of projection cells, interneurons, excitatory and inhibitory neurotransmission, and various molecular machineries. Our data demonstrate that CaMKIIα and GAD1 cell populations are distinct in avian pallium, and that they mirror several physiological features of excitatory and inhibitory neocortical neurons. These observations strongly suggest that 1) this set of shared features are core physiological properties that are critical for pallial function, and 2) a shared ancestral origin of cell types and circuit elements constrains the evolution of divergent physiological phenotypes between bird and mammalian pallium.

*In vitro*, mammalian neocortical excitatory neurons can be distinguished from inhibitory interneurons by the electrical phenotype of continuous accommodation to depolarizing current injections^71^. In the present study in finches, CaMKIIα neurons exhibit very strong adaptation by firing 1-2 phasic action potentials throughout the current step protocol, whereas GAD1 neurons exhibit a non-accomodating electrical phenotype. *In vivo* physiological recordings showed that CaMKIIα and GAD1 neurons can also be distinguished as broad-spiking and narrow-spiking, respectively, as in mammalian neocortex. We extend previous findings in songbirds demonstrating that auditory coding roles of broad-spiking and narrow-spiking neurons show strong parallels with the role of excitatory neurons and inhibitory interneurons in mammalian auditory cortex^7,28,35–37,39,40^, and attribute those properties to CaMKIIα vs. GAD1 neurons specifically. Additionally, our findings suggest that in avian auditory pallium, GAD1 neurons may tune stimulus selectivity of excitatory neurons, a functional outcome predicted directly from recent data in rodent auditory cortex^61^. These data strongly imply that in birds as in mammals, the molecular identity of excitatory vs. inhibitory neurons in pallium is tied to a fundamental set of physiological and sensory coding properties. The relationship between molecular identities and physiological roles exist irrespective of their anatomical organization, such as the denser, clustered organization of pallial neurons in birds compared to less dense, layered mammalian neocortex^72,73^.

Our data also highlight similarities between CaMKIIα and GAD1 cells in birds and subclasses of excitatory and inhibitory cell types in mammalian pallium. CaMKIIα neurons were largely non-pyramidal in morphology, and previous work has not identified projections from NCM that exit pallium (though see ^74^). NCM instead makes reciprocal connections with other pallial domains, including the mesopallium, a large territory with neurons that recent developmental and genetic data demonstrate to have similar molecular markers to mammalian intra-telencephalic (IT) cells^12,22^. A majority of CaMKIIα cells in this study could conceivably represent a class of conserved IT pallial cells^75^, though this requires further study. Our GAD1 population exhibits a fair degree of variance in auditory coding properties (Fig. 3), suggesting we may be capturing multiple subclasses of inhibitory interneurons with this viral construct. Previous work has demonstrated there is immunoreactivity in NCM and other pallial regions for parvalbumin (PV) as well as calbindin, and that these cells invade pallium during development in a tangential migration similar to mammals^15,72,76,77^. In the present study we observed that GAD1-positive cells (and mDlx cells) co-localize with PV (Fig. 1D,E). The physiological roles of GAD1 neurons in NCM that we observed, including a fast-spiking phenotype, faster response latencies, broader selectivity for auditory stimuli, and control over gamma oscillations, are consistent with the role of PV cells in mammalian neocortex^53,78,79^. Future studies matching molecular identities to physiological properties will be necessary to determine the extent to which subclasses of pallial cell types have diverged or retained intrinsic physiological properties across amniotes. Additionally, future studies of projection patterns within and outside the telencephalon, combined with dendritic and axonal morphology of CaMKIIα and GAD1 cell types, will further clarify the distinctions and similarities between avian and mammalian pallial cell types.

At the network level, we find several similarities in birds compared to mammals in the organization of cell types. Driving GAD1 and mDlx neurons induces local suppression and synchronizes networks of narrow-spiking cells at the level of single units and the broad-scale gamma oscillations in the surrounding pallium. Suppressing GAD1 neurons has the opposite effect on pallial synchrony. Suppressing GAD1 neurons also altered stimulus selectivity of broad-spiking single units, suggesting inhibitory control of downstream excitatory sensory coding. However, contrary to prediction, suppression of GAD1 cells did not alter firing rate of broad-spiking units. This suggests a species-level difference in the role of interneurons in constraining firing rate, but more precise cell-type manipulations could reveal specific effects of PV+ SOM+ or VIP+ neurons on firing rate that are obfuscated in the present study by simultaneous activation of multiple interneuron types with opposing network roles ^80–82^. Pallial circuits in birds may therefore exhibit further similarities to mammalian cortical microcircuits, including feedforward suppression^56,57^, though our data do not preclude alternative circuit mechanisms for local signal processing.

Our current findings highlight both similarities and differences that reveal core intrinsic and network features of excitatory neurons and inhibitory neurons that either evolved independently or are conserved features of pallium in amniotes^83^. In either case, the retention- or convergent evolution-of certain core physiological and network features of pallial cell types in birds and mammals clarifies candidate features of pallium essential for advanced cognition.

## Materials and Methods

### Animals

Adult zebra finches used in this study were housed in unisex aviaries prior to experiments under a photoperiod of 14-h light:10-h dark. Birds were provided with food and water ad libitum, as well as several forms of weekly dietary enrichment (e.g., egg food, fresh millet branches, cuttlebone). All procedures and protocols adhered to the guidelines of the National Institutes of Health Guide for the Care and Use of Laboratory Animals, and were approved by the University of Massachusetts, Amherst Institutional Animal Care and Use Committee.

### Virus injection surgeries

4-6 weeks prior to optogenetic experiments, we injected viruses bilaterally into the caudomedial nidopallium (NCM; the avian auditory association pallium) of male and female zebra finches. Birds were fasted ~30 min prior to surgery, anesthetized with 2% isoflurane in 2 L/min O2, fixed to a custom stereotax (Herb Adams Engineering) equipped with a heating pad (DC Neurocraft) at a 45° head angle, and maintained on 1.5% isoflurane, 1 L/min O2 for the duration of the surgery. Points overlying NCM were marked by scoring the skull lateral and rostral to our coordinates (i.e., a crosshair), and a craniotomy exposed the brain surface (NCM coordinates = 1.1 mm rostral, 0.7 mm lateral of stereotaxic zero, defined as the caudal edge of the bifurcation of the midsaggital sinus). A glass pipette (tip diameter: 20-50 μm) filled with mineral oil was loaded with virus (for constructs and titers see below), attached to a Nanoject (Drummond Scientific Company, Broomall, PA), and lowered into the brain at a depth of 1.5 mm from the surface. We injected 625 nL of virus, 2 nL/sec, 125 nL/cycle, 5 cycles, with a 60 sec wait between cycles. Following injections, we waited 10 min before slowly retracting the injection pipette from the brain. After injections were made in both hemispheres, crainiotomies were filled using Kwik-Cast (World Precision Instruments, Sarasota, FL), and the scalp was fixed around the crainiotomy using cyanoacrylate adhesive.

The viruses and titers used in the present study included: CaMKIIα, pAAV9-CaMKIIα-hChR2(E123A)-EYFP (titer: 1×10^13^ viral genomes/mL; Addgene: 35505); GAD1, pAAV9-hGAD1-GLAD-NLS-CRE (titer: 1.25×10^12^ viral genomes/mL; made in-house), pAAV9-mDlx-ChR2(H134R)-EYFP (titer: 1.58×10^12^ viral genomes/mL; made in-house), AAV9 pCAG-FLEX-tdTomato-WPRE (titer: 1.5×10^12^ viral genomes/mL; Addgene: 51503).

### Immunofluorescence

A subset of birds (N = 3; 2 males, 1 female) given injections with viruses targeting both CaMKIIα and GAD1 promoters were transcardially perfused with 4 % paraformaldehyde 4-6 weeks following surgery. Brains were extracted, post-fixed overnight in 4 % paraformaldehyde, then dehydrated in 30 % sucrose in 0.1 M phosphate-buffered saline (PBS). After dehydration, brains were placed in 2 x 2 x 2 inch plastic blocks filled with cryo-embedding compound (Ted Pella Inc., Redding, CA). Brains were sectioned coronally at 40 microns and sections were stored in cryoprotectant solution (30 % sucrose, 1 % polyvinylpyrrolidone, 30 % ethylene glycol, in 0.1 M phosphate buffer) at −20 °C.

We labeled tissue sections for aromatase, a reliable marker for NCM compared to neighboring regions^72,84^. At all times during this procedure tissue was processed in a shaded room away from direct light sources. Tissue sections were washed 5 times in 0.1 M PBS, 3 times in 0.1 M phosphate-buffered saline with 0.3% triton (0.3 % PBT), blocked using 10 % normal goat serum, and incubated at 4 °C for two days in rabbit anti-aromatase diluted 1:2000 in blocking serum (aromatase antibody provided as a generous gift from Dr. Colin Saldanha). In N = 1 bird, tissue was simultaneously incubated in mouse anti-NeuN (Millipore, Danvers, MA; RRID: AB_2298772) diluted 1:2000 in blocking serum.

Sections were then washed 3 times in 0.1 % PBT and incubated in secondary antibodies diluted 1:500 in 0.3 % PBT. Secondaries used were goat anti-mouse Alexa 405 (Abcam, Cambridge, MA; for N = 1 bird, tissue that was incubated in mouse anti-NeuN above) and goat anti-rabbit Alexa 647 (Invitrogen, Waltham, MA; for N = 3 birds). After 3 more washes in 0.1 % PBT, sections were mounted onto slides, and coverslipped using Prolong antifade mounting medium with DAPI stain in the 405 channel (for bird with tissue incubated in mouse anti-NeuN, Prolong mounting medium contained no DAPI; Invitrogen, Waltham, MA). Slides were dried overnight at room temperature, and stored at 4 °C until imaging.

Sections were imaged on a confocal microscope (A1SP; Nikon, Tokyo, Japan) at the UMass light microscopy core facility. Images were acquired using NIS-Elements software (RRID: SCR_002776). We determined gain and laser intensity separately for each tissue section to minimize background fluorescence. Injection sites were localized within NCM by dense fluorescence in the aromatase-positive field of NCM. To ensure we were only capturing images of transduced cells within NCM, pictures of ventral and dorsal NCM were taken only within the aromatase-positive population of the region dorso-medial to the medial part of the dorsal arcopallial lamina.

Images of tissue sections containing NCM were first taken at 10X to serve as a reference for higher magnification imaging. All cells within an image in NCM were taken at 60X using z-stacks of 1-2 μm through the entire thickness of the tissue section.

#### Analysis

For all confocal images we quantified the number of EYFP-positive cells, the number of tdTomato-positive cells, and the number of cells expressing both fluorophores (reflecting transduction of the CaMKIIα and GAD1 promoter, or both, respectively) The total number of transduced cells across all observed slices (n = 122) were tallied. Cells were counted only when positive staining was associated with DAPI or NeuN. In slices that were stained with antibody targeting NeuN, no labeled cells were observed that did not co-localize with NeuN labeling.

### *In vitro* neuronal response properties

Birds were rapidly decapitated and dissected, then the caudal telencephalon was bisected on a petri dish immersed in wet ice and each hemisphere was sectioned at 250 microns on a vibratome. A glycerin-based external cutting solution (substituting for NaCl & sucrose) was used for sectioning to improve slice survival time ^85^, containing (in mM): 222 glycerin, 25 NaHCO_3_, 2.5 KCl, 1.25 NaH_2_PO_4_, 0.5 CaCl_2_, 3 MgCl_2_•6H_2_O, 25 dextrose, 0.4 ascorbic acid, 2 sodium pyruvate, 3 myo-inositol; 310 mOsm/kg H_2_O; pH 7.4 when saturated with 95% O_2_/5% CO_2_. Following sectioning, the slices were warmed to ~ 40 °C in external solution containing (in mM): 111 NaCl, 25 NaHCO_3_, 2.5 KCl, 1.25 NaH_2_PO_4_, 2 CaCl_2_, 1 MgCl_2_•6H_2_O, 25 dextrose, 0.4 ascorbic acid, 2 sodium pyruvate, 3 myoinositol; 310 mOsm/kg H_2_O; pH 7.4 when saturated with 95% O_2_/5% CO_2_. Our potassium-based internal solution contained (in mM): 2.4 potassium gluconate, 0.4 KCl, 0.002 CaCl_2_, 0.1 HEPES, 0.1 EGTA, 0.06 Mg-ATP, 0.01 Na-GTP, 0.4 C4H8N3O5PNa2•4H2O; 290-305 mOsm/kg H2O; pH 7.4.

Tissue was imaged using a charge-coupled camera (QIClick; QImaging) mounted to a fixed stage microscope (Eclipse FN1; Nikon) that was equipped with a water emersion objective (CFI Fluor; 40X; NA = 0.8; WD = 2.0 mm; Nikon). Glass pipettes were pulled from borosilicate glass capillary tubes (1B150F-4; World Precision Instruments) using a two-stage, vertical puller (PC-10; Narishige International USA). Positive pressure was applied to each pipette while moving through aCSF and tissue. Liquid junction potential was automatically subtracted. Pipettes had a tip resistance of 4-8 MΩ when backfilled with internal solution, which routinely included neurobiotin tracer (Vector Laboratories) for identification post-recording. Once a whole-cell configuration was successfully achieved, series resistance and slow capacitive transients were counterbalanced. Each recording configuration was allowed to stabilize for a minimum of five min before beginning protocols ^86^. Experimental data were acquired at 40 kHz using multi-channel acquisition software (PATCHMASTER; HEKA Elektronik), digitized using a patch clamp amplifier (EPC 10 USB; HEKA Elektronik), then exported to Igor Pro (WaveMetrics) and processed using a 1 kHz low-pass filter.

Post-recording, tissue sections were drop fixed in 4% Paraformaldehyde dissolved in 0.025 M phosphate buffer (PB), then stored at −20 °C in a cryoprotectant solution composed of 30% sucrose, 30% ethylene glycol, and 1% polyvinylpyrrolidone in 0.1 M phosphate buffer (PB).

#### Analysis

Recordings were made from N = 5 birds (N = 3 males, N = 2 females), n = 18 cells transduced with CaMKIIα-ChR2 and N = 4 birds N= 2 males, N = 2 females), n = 11 cells transduced with GAD1-ChR2. Steady-state voltage (SSV), action potential kinetics, and action potential width (duration at 50% of action potential depolarization peak from resting membrane potential) were each calculated using custom scripts written in Python.

### Anesthetized extracellular recording

Extracellular recording was performed *in vivo* under urethane anesthesia as in previous studies ^87^. On the day of recording, birds were injected in the pectoral muscle with 90-120 μL 20% urethane (30 μL every 45 min; specific amount depended on the mass of the bird). Anesthetized birds were fixed to a custom stereotaxic apparatus as above and a stainless steel headpost was fixed to the head using acrylic cement. Birds were then moved to a sound-attenuation booth (Industrial Acoustics) on an air table (TMC, Peabody, MA) for optotagging and extracellular recording experiments. In the booth birds were fixed to a custom stereotax (Herb Adams Engineering) at a 45° head angle using the attached headpost and the craniotomy above NCM was exposed. We recorded preferentially from the left hemisphere but both hemispheres are represented in our dataset.

An optrode consisting of a single tungsten electrode (A-M Systems, Sequim, WA) epoxied to an optic fiber (diameter: ~200 μm; Thor Labs, Newton, NJ; distance between electrode and fiber: 400-600 μm) was lowered into the brain at the NCM coordinates specified above. The optrode was calibrated by inserting the optrode into a light meter (Thor Labs, Newton, NJ) in a dark room, adjusting laser power and obtaining a calibration curve of laser setting to output wattage. To elicit ChR2-induced spike activity locally around our electrode we used a blue laser (447 nm) with an output wattage of ~2 mW. Recordings were made between 1.2 and 1.8 mm ventral to the brain surface to search for characteristic NCM baseline and sound-evoked activity, as well as light-evoked activity. Light stimuli consisted of either 25 msec pulses or 100 ms pulses, and pulses were separated by 4-10 sec. Following light-only trials where light-evoked units were identified (see below), we immediately ran auditory trials in the same location. Auditory trials consisted of seven stimuli: 6 conspecific songs and white noise, all normalized to ~70 decibels. Stimuli were randomly presented during trials and each stimulus was presented 15 times. At a given recording site, we randomly presented 3 of 6 conspecific songs and white noise (i.e., 4 stimuli total). Interstimulus interval was 10+2 sec. Recordings were amplified, bandpass filtered (300 to 5000 Hz; A-M Systems), and digitized at 16.67 kHz (Micro 1401, Spike2 software; Cambridge Electronic Design). Following recordings, birds were transcardially perfused and brains were extracted as above for anatomical confirmation of optrode sites.

#### Analysis

Data were processed in Spike2 (version 7.04) to identify light-evoked units. Recordings from light-only trials were thresholded above the noise band and peristimulus histograms and raster plots were generated to examine multiunit responses to light stimulation. We only performed auditory playback trials if there was a peak in the histogram during the light stimulus that was consistent across trials (as shown by the raster plot; Fig. 3A). Recordings were made from N = 3 birds (1 male, 2 females), n = 22 cells transduced with CaMKIIα-ChR2 and N = 3 birds (3 females), n = 16 cells transduced with GAD1-ChR2.

Templates for single units in light-only trials were isolated in Spike2 by their waveform characteristics and filtered so that no units had an interspike interval > 1 ms as in previous studies ^87^. Individual spikes were assigned to generated templates with an accuracy range of 60%-100%. Principal component analyses were used to confirm well-isolated units (i.e., non-overlapping clusters in 3-dimensional space). We could reliably obtain 2-4 units from each recording site. We used isolated single unit waveforms from light-only trials to sort multiunit activity in their paired auditory trials. At this point we again checked that none of the units had interspike intervals > 1 ms and that principal component analyses still demonstrated non-overlapping clusters.

For waveform analyses, we compared physiological properties of light-responsive cells between birds with virus targeting the CaMKIIα promoter and birds with virus targeting the GAD1 promoter. We calculated action potential width as the time between the first (depolarized) and second (hyperpolarized) peaks of the average waveform. Spike quarter-width was calculated as the width of the waveform at a quarter of the height of the action potential.

Z scores were calculated for each single unit’s response to conspecific song and white noise to provide a normalized change in firing during stimulus presentation (*S*) compared to baseline (*B*), using the following formula as in previous studies ^88^:

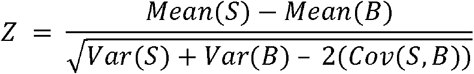

Z scores for conspecific songs for each single unit are reported as average single unit responses across conspecific stimuli presented.

We calculated single unit response latency to white noise by adapting a previously described method^40^. We calculated the mean and standard deviation of the baseline period (2 s before stimulus onset). Then, peri-stimulus time histograms were created for each single unit’s response to white noise, divided into 5 ms bins and smoothed with a 5-point boxcar filter. We identified the first bin within 400 ms of white noise onset in which firing rate passed a threshold of 3 standard deviations from the baseline mean.

To examine auditory coding properties of isolated single units, we used a custom pattern classifier ^39,89^. The classifier uses timing accuracy of stimulus-evoked firing responses of single units to discriminate amongst the different stimulus types, providing a measure of how well stimuli can be distinguished by consistency of evoked firing across trials.

Specifically, for each single unit the classifier pseudorandomly picked one response per stimulus to serve as templates. The classifier compared the selected templates with all other stimulus-evoked responses of that single unit. Based on values of a distance metric (detailed below) between responses and the templates, the algorithm would assign a template identity (e.g. conspecific song 1) to the template that was most similar to a given response. The classifier repeated this procedure 1000 times and then determined the mean accuracy of assigning stimulus-evoked responses to their correct associated stimuli.

For the timing accuracy measure, data were convolved with Gaussian filters prior to template comparison. The optimal standard deviation of the filter for each cell was picked based on which value gave the highest accuracy (values that were used: 1, 2, 4, 8, 16, 32, 64, 128,

256 ms). Comparisons between templates and responses used the R_corr_ distance metric ^39,90^.

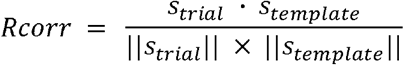

Where s represents the vectors of the trial and the template responses after filtering, dot multiplied and divided by the product of their lengths. For the timing accuracy measure the classifier matched responses (i.e., trials) to templates based on which template yielded the highest R_corr_ value.

To statistically test whether the accuracy of the classifier was greater than random chance, we generated confusion matrices and used a trial shuffling approach (modified from ^39,89^). Classifier assignments of stimulus-evoked responses in the confusion matrix were shuffled and randomly assigned to stimuli 1000 times. The distribution of the classifier accuracies across the 1000 runs was compared to the randomly shuffled assignments. Accuracies were considered significantly greater than random when Cohen’s d was > 0.2 ^39,91^. All single unit accuracies in this study had a Cohen’s d > 0.2. Analyses for conspecific song responses are presented as single unit responses averaged across conspecific stimuli presented to a given unit.

Temporal stimulus selectivity was measured by comparing the mean R_corr_ values across stimuli repetitions (spike-timing correlation) among the 3 different conspecific stimuli. Specifically, for each single unit, the 3 mean R_corr_ values (Rx, Ry, Rz) were used to generate a 3-dimensional vector 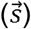. Then, a second vector was generated such that vector magnitude was identical to 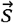 but R_corr_ values for stimuli were equalized in the 3D-space (Rx = Ry = Rz), i.e. the equal-selectivity vector 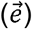. The angle (in radians) between each single unit’s 3D vector 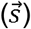 and its associated equal-selectivity vector 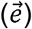 was calculated using the arccosine distance metric: 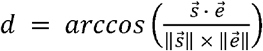, where values close to 0 reflect non-selectivity and larger values reflect greater selectivity.

### Optodrive awake recordings

For larger scale recordings, custom optodrives were made by coupling a fiber optic (200 μm diameter) to 8 tetrodes arranged in a single bundle (customized from EIB Open-Ephys drive; Open Ephys Inc., MA). Tetrodes were made from 12.5 μm insulated NiCr wires (Sandvik, Sandviken, Sweden). The optic fiber was placed ~600 μm above the wires. The horizontal distance between the fiber and wire bundle was ~200 μm.

Virally transduced birds were anesthetized with isoflurane, after which bilateral craniotomies were made over NCM (see above for coordinates; sealed with Kwik-Cast) and headposts were secured to the dorsomedial rostral skull with acrylic cement. 2-4 days later, birds were restrained and head-fixed. Optodrives were lowered into NCM 1.5-2.0 mm ventral of the brain surface. After finding a site with characteristic NCM baseline and sound-evoked activity, the optodrive was allowed to stabilize in the brain for ~1 h. Following stabilization, a blue (447 nm) or green (532 nm) laser was used to test network responses to manipulation of transduced cells. Recordings were obtained using the Open-Ephys GUI ^92^, and amplified and digitized at 30 kHz using Intan Technologies amplifier and evaluation board (RHD2000; Intan Technologies, Los Angeles, CA). Laser delivery was controlled by custom MatLab (MathWorks, Natick, MA) scripts and an Arduino Uno integrated with the evaluation board.

For the GAD-1 archaerhodopsin experiment, after testing network responses to manipulation of transduced cells by the green laser, we tested how inhibition of GAD-1 cells would affect functional auditory response properties of the network. We randomized playback of 4 conspecific songs. Each song was heard 10 times total, 5 times without the green laser, and 5 times with the green laser turned on specifically for the duration of the stimulus. This resulted in 20 playback trials with laser turned off (and songs randomized within these 20 trials such that each song was played 5 times), followed by 20 playback trials with laser turned on only during the duration of each song playback (and songs randomized within these 20 trials such that each song was played 5 times). We allowed 5 seconds of silence in between playback of different stimuli.

#### Analysis

Optodrive single unit sorting was done with Kilosort ^93^. Sorting results were manually curated and only well-isolated units (high signal-to-noise ratio; low contamination; good segregation in waveform principal component analysis space; low frequency in violations of refractory period) were used for these analyses. Data were common median filtered and 300 Hz high-pass filtered.

Sample sizes for optodrive experiments were: mDlx-ChR2: N = 1 female; n = 36 single units. GAD1-ChR2: N = 1 male; n = 30 single units. GAD1-archaerhodopsin: N = 1 male; n = 52 single units. Recordings were made from both left and right hemispheres.

For local field potential (LFP) analyses, raw traces were 150 Hz low-pass filtered and local field potential power spectra were generated in Python (smoothed and imported using Neo ^94^ and Elephant libraries; Welch’s power spectrum frequency resolution at 1 Hz).

We also calculated stimulus firing rate in Hertz and auditory selectivity (as described above in the analysis section for anaesthetized optrode recording) for the GAD1-archaerhodopsin experiment to compare responses to conspecific song when laser was off versus when the laser was on.

### Statistics

Statistical analyses included two-tailed t-tests and nonparametric statistics (Mann Whitney U tests) when log_10_ transformations did not correct violations of parametric assumptions. Bonferroni corrections were used to correct for multiple comparisons. Wilcoxon signed rank tests were used to test paired responses to song in the same cells when laser was off compared to on. Chi-square tests were used to test whether the number of cells that increased or decreased firing rates and auditory selectivity was greater than expected by chance.

## Data availability

Data from this study will be made available on Dryad.

## Acknowledgements

We thank M. Fernandez-Peters, K. Schroeder, C. Healey, and P. Katz for feedback on manuscript, and H. Boyd, N. Ambrosio, and V. Ivan for assistance with data collection.

## Author contributions

J.S., Y.Y.-S., and L.R.-H. designed experiments; Y.M. designed viruses for experiments; J.S., M.M.-L., G.S., and L.R.-H. performed experiments, wrote code, and analyzed data. J.S. and L.R.-H. wrote the paper.

## Supplementary Information

Figures S1-S7

## References

1. Briscoe, S. D. & Ragsdale, C. W. Evolution of the Chordate Telencephalon. Current Biology vol. 29 R647–R662 (2019).

2. Kaas, J. H. The evolution of brains from early mammals to humans. Wiley Interdiscip. Rev. Cogn. Sci. 4, 33–45 (2013).

3. Mountcastle, V. B. The columnar organization of the neocortex. Brain vol. 120 701–722 (1997).

4. Reiner, A. et al. Revised nomenclature for avian telencephalon and some related brainstem nuclei. J. Comp. Neurol. 473, 377–414 (2004).

5. Jarvis, E. et al. Avian brains and a new understanding of vertebrate brain evolution. Nature Reviews Neuroscience vol. 6 151–159 (2005).

6. Wang, Y., Brzozowska-Prechtl, A. & Karten, H. J. Laminar and columnar auditory cortex in avian brain. Proc. Natl. Acad. Sci. 107, 12676–12681 (2010).

7. Calabrese, A. & Woolley, S. M. N. Coding principles of the canonical cortical microcircuit in the avian brain. Proc. Natl. Acad. Sci. 112, 3517–3522 (2015).

8. Stacho, M. et al. A cortex-like canonical circuit in the avian forebrain. Science (80-.). 369, (2020).

9. Shimizu, T., Cox, K. & Karten, H. J. Intratelencephalic projections of the visual wulst in pigeons (Columba livia). J. Comp. Neurol. 359, 551–572 (1995).

10. Güntürkün, O. & Bugnyar, T. Cognition without Cortex. (2016) doi:10.1016/j.tics.2016.02.001.

11. Suryanarayana, S. M., Robertson, B., Wallén, P. & Grillner, S. The Lamprey Pallium Provides a Blueprint of the Mammalian Layered Cortex. Curr. Biol. 27, 3264–3277.e5 (2017).

12. Briscoe, S. D., Albertin, C. B., Rowell, J. J. & Ragsdale, C. W. Neocortical Association Cell Types in the Forebrain of Birds and Alligators. Curr. Biol. 28, 686–696.e6 (2018).

13. Tosches, M. A. et al. Evolution of pallium, hippocampus, and cortical cell types revealed by single-cell transcriptomics in reptiles. Science (80-.). 360, 881–888 (2018).

14. Suzuki, I. K. & Hirata, T. Neocortical neurogenesis is not really ‘neo’: A new evolutionary model derived from a comparative study of chick pallial development. Development Growth and Differentiation vol. 55 173–187 (2013).

15. Cobos, I., Puelles, L., Martínez, S., Hernandez, M. & Juan, S. The Avian Telencephalic Subpallium Originates Inhibitory Neurons That Invade Tangentially the Pallium (Dorsal Ventricular Ridge and Cortical Areas). (2001) doi:10.1006/dbio.2001.0422.

16. Puelles, L. et al. The pallium in reptiles and birds in the light of the updated tetrapartite pallium model. in Evolution of Nervous Systems (ed. Kaas, J.) 519–555 (Academic Press, 2017).

17. Jarvis, E. D. et al. Global view of the functional molecular organization of the avian cerebrum: Mirror images and functional columns. J. Comp. Neurol. 521, 3614–3665 (2013).

18. Desfilis, E., Abellán, A., Sentandreu, V. & Medina, L. Expression of regulatory genes in the embryonic brain of a lizard and implications for understanding pallial organization and evolution. J. Comp. Neurol. 526, 166–202 (2018).

19. Butler, A. B., Reiner, A. & Karten, H. J. Evolution of the amniote pallium and the origins of mammalian neocortex. Ann. N. Y. Acad. Sci. 1225, 14–27 (2011).

20. Martin Wild, J., Karten, H. J. & Frost, B. J. Connections of the auditory forebrain in the pigeon (columba livia). J. Comp. Neurol. 337, 32–62 (1993).

21. Zeier, H. & Karten, H. J. The Archistriatum of the Pigeon: Organization of Afferent and Efferent Connections. Brain Res. 31, 313–326 (1971).

22. Vates, G. E., Broome, B. M., Mello, C. V. & Nottebohm, F. Auditory pathways of caudal telencephalon and their relation to the song system of adult male zebra finches (Taenopygia guttata). J. Comp. Neurol. 366, 613–642 (1996).

23. Karten, H. J. & Shimizu, T. The Origins of Neocortex: Connections and Lamination as Distinct Events in Evolution. J. Cogn. Neurosci. 1, 291–301 (1989).

24. Chew, S. J., Vicario, D. S. & Nottebohm, F. A large-capacity memory system that recognizes the calls and songs of individual birds. Proc. Natl. Acad. Sci. U. S. A. 93, 1950–1955 (1996).

25. Phan, M. L., Pytte, C. L. & Vicario, D. S. Early auditory experience generates long-lasting memories that may subserve vocal learning in songbirds. Proc. Natl. Acad. Sci. U. S. A. 103, 1088–1093 (2006).

26. Gobes, S. M. H. & Bolhuis, J. J. Birdsong Memory: A Neural Dissociation between Song Recognition and Production. Curr. Biol. 17, 789–793 (2007).

27. Gentner, T. Q. & Margoliash, D. Neuronal populations and single cells representing learned auditory objects. Nature 424, 669–674 (2003).

28. Yanagihara, S. & Yazaki-Sugiyama, Y. Auditory experience-dependent cortical circuit shaping for memory formation in bird song learning. Nat. Commun. 7, 1–11 (2016).

29. Sen, K., Theunissen, F. E. & Doupe, A. J. Feature Analysis of Natural Sounds in the Songbird Auditory Forebrain. J. Neurophysiol. 86, 1445–1458 (2001).

30. Shimizu, T., Patton, T. B. & Husband, S. A. Avian visual behavior and the organization of the telencephalon. Brain. Behav. Evol. 75, 204–217 (2010).

31. Ditz, H. M. & Nieder, A. Neurons selective to the number of visual items in the corvid songbird endbrain. Proc. Natl. Acad. Sci. U. S. A. 112, 7827–7832 (2015).

32. Moll, F. W. & Nieder, A. Cross-modal associative mnemonic signals in crow endbrain neurons. Curr. Biol. 25, 2196–2201 (2015).

33. Veit, L. & Nieder, A. Abstract rule neurons in the endbrain support intelligent behaviour in corvid songbirds. Nat. Commun. 4, 1–11 (2013).

34. Dugas-Ford, J., Rowell, J. J. & Ragsdale, C. W. Cell-type homologies and the origins of the neocortex. Proc. Natl. Acad. Sci. 109, 16974–16979 (2012).

35. Meliza, C. D. & Margoliash, D. Emergence of selectivity and tolerance in the avian auditory cortex. J. Neurosci. 32, 15158–68 (2012).

36. Bottjer, S. W., Ronald, A. A. & Kaye, T. Response properties of single neurons in higher level auditory cortex of adult songbirds. J Neuro-physiol 121, 218–237 (2019).

37. Schneider, D. M. & Woolley, S. M. N. Sparse and Background-Invariant Coding of Vocalizations in Auditory Scenes. Neuron 79, 141–152 (2013).

38. Moseley, D. L., Joshi, N. R., Prather, J. F., Podos, J. & Remage-Healey, L. A neuronal signature of accurate imitative learning in wild-caught songbirds (swamp sparrows, Melospiza georgiana). Sci. Rep. 7, 17320 (2017).

39. Krentzel, A. A., Macedo-Lima, M., Ikeda, M. Z. & Remage-Healey, L. A Membrane G-Protein-Coupled Estrogen Receptor Is Necessary but Not Sufficient for Sex Differences in Zebra Finch Auditory Coding. Endocrinology 159, 1360–1376 (2018).

40. Ono, S., Okanoya, K. & Seki, Y. Hierarchical emergence of sequence sensitivity in the songbird auditory forebrain. J. Comp. Physiol. A Neuroethol. Sensory, Neural, Behav. Physiol. 202, 163–183 (2016).

41. Mooney, R. & Prather, J. F. The HVC microcircuit: The synaptic basis for interactions between song motor and vocal plasticity pathways. J. Neurosci. 25, 1952–1964 (2005).

42. Rauske, P. L., Shea, S. D. & Margoliash, D. State and Neuronal Class-Dependent Reconfiguration in the Avian Song System. J. Neurophysiol. 89, 1688–1701 (2003).

43. Pfeffer, C. K., Xue, M., He, M., Huang, Z. J. & Scanziani, M. Inhibition of inhibition in visual cortex: The logic of connections between molecularly distinct interneurons. Nat. Neurosci. 16, 1068–1076 (2013).

44. Lee, S. H. et al. Activation of specific interneurons improves V1 feature selectivity and visual perception. Nature 488, 379–383 (2012).

45. Dimidschstein, J. et al. A viral strategy for targeting and manipulating interneurons across vertebrate species. Nat. Neurosci. 19, 1743–1749 (2016).

46. Meyer, H. S. et al. Inhibitory interneurons in a cortical column form hot zones of inhibition in layers 2 and 5A. Proc. Natl. Acad. Sci. U. S. A. 108, 16807–16812 (2011).

47. Kawaguchi, Y. Groupings of nonpyramidal and pyramidal cells with specific physiological and morphological characteristics in rat frontal cortex. Artic. J. Neurophysiol. 69, (1993).

48. McCormick, D. A., Connors, B. W., Lighthall, J. W. & Prince, D. A. Comparative electrophysiology of pyramidal and sparsely spiny stellate neurons of the neocortex. J. Neurophysiol. 54, 782–806 (1985).

49. Lo, F.-S., Akkentli, F., Tsytsarev, V. & Erzurumlu, R. S. Functional significance of cortical NMDA receptors in somatosensory information processing. J. Neurophysiol. 110, 2627–2636 (2013).

50. Gentet, L. J. et al. Unique functional properties of somatostatin-expressing GABAergic neurons in mouse barrel cortex. Nat. Neurosci. 15, 607–612 (2012).

51. Suzuki, N. & Bekkers, J. M. Microcircuits mediating feedforward and feedback synaptic inhibition in the piriform cortex. J. Neurosci. 32, 919–931 (2012).

52. Lou, S. et al. Genetically Targeted All-Optical Electrophysiology with a Transgenic Cre-Dependent Optopatch Mouse. J. Neurosci. 36, 11059–11073 (2016).

53. Moore, A. K. & Wehr, M. Parvalbumin-expressing inhibitory interneurons in auditory cortex are well-tuned for frequency. J. Neurosci. 33, 13713–13723 (2013).

54. Atencio, C. A. & Schreiner, C. E. Spectrotemporal processing differences between auditory cortical fast-spiking and regular-spiking neurons. J. Neurosci. 28, 3897–3910 (2008).

55. Sakata, S. & Harris, K. D. Laminar Structure of Spontaneous and Sensory-Evoked Population Activity in Auditory Cortex. Neuron 64, 404–418 (2009).

56. Wehr, M. & Zador, A. M. Balanced inhibition underlies tuning and sharpens spike timing in auditory cortex. Nature 426, 442–446 (2003).

57. Cruikshank, S. J., Lewis, T. J. & Connors, B. W. Synaptic basis for intense thalamocortical activation of feedforward inhibitory cells in neocortex. Nat. Neurosci. 10, 462–468 (2007).

58. Cardin, J. A. Inhibitory Interneurons Regulate Temporal Precision and Correlations in Cortical Circuits. Trends in Neurosciences vol. 41 689–700 (2018).

59. Sohal, V. S., Zhang, F., Yizhar, O. & Deisseroth, K. Parvalbumin neurons and gamma rhythms enhance cortical circuit performance. Nature 459, 698–702 (2009).

60. Isaacson, J. S. & Scanziani, M. How inhibition shapes cortical activity. Neuron vol. 72 231–243 (2011).

61. Moore, A. K., Weible, A. P., Balmer, T. S., Trussell, L. O. & Wehr, M. Rapid Rebalancing of Excitation and Inhibition by Cortical Circuitry. Neuron 97, 1341–1355.e6 (2018).

62. Klug, J. R. et al. Genetic Inhibition of CaMKII in Dorsal Striatal Medium Spiny Neurons Reduces Functional Excitatory Synapses and Enhances Intrinsic Excitability. PLoS One 7, (2012).

63. Gerfen, C. R. Synaptic organization of the striatum. J. Electron Microsc. Tech. 10, 265–281 (1988).

64. Tepper, J. M. et al. Heterogeneity and Diversity of Striatal GABAergic Interneurons: Update 2018. Front. Neuroanat. 12, 91 (2018).

65. Siemian, J. N., Sarsfield, S. & Aponte, Y. Glutamatergic fast-spiking parvalbumin neurons in the lateral hypothalamus: Electrophysiological properties to behavior. Physiology and Behavior vol. 221 112912 (2020).

66. Poulin, J. F., Castonguay-Lebel, Z., Laforest, S. & Drolet, G. Enkephalin co-expression with classic neurotransmitters in the amygdaloid complex of the rat. J. Comp. Neurol. 506, 943–959 (2008).

67. Poulin, J. F., Arbour, D., Laforest, S. & Drolet, G. Neuroanatomical characterization of endogenous opioids in the bed nucleus of the stria terminalis. Prog. Neuro-Psychopharmacology Biol. Psychiatry 33, 1356–1365 (2009).

68. Nair-Roberts, R. G. et al. Stereological estimates of dopaminergic, GABAergic and glutamatergic neurons in the ventral tegmental area, substantia nigra and retrorubral field in the rat. Neuroscience 152, 1024–1031 (2008).

69. Yoo, J. H. et al. Ventral tegmental area glutamate neurons co-release GABA and promote positive reinforcement. Nat. Commun. 7, 1–13 (2016).

70. Root, D. H. et al. Selective Brain Distribution and Distinctive Synaptic Architecture of Dual Glutamatergic-GABAergic Neurons. Cell Rep. 23, 3465–3479 (2018).

71. Markram, H. et al. Reconstruction and Simulation of Neocortical Microcircuitry. Cell 163, 456–492 (2015).

72. Ikeda, M. Z., Krentzel, A. A., Oliver, T. J., Scarpa, G. B. & Remage-Healey, L. Clustered organization and region-specific identities of estrogen-producing neurons in the forebrain of Zebra Finches (Taeniopygia guttata). J. Comp. Neurol. 525, 3636–3652 (2017).

73. Olkowicz, S. et al. Birds have primate-like numbers of neurons in the forebrain. Proc. Natl. Acad. Sci. U. S. A. 113, 7255–7260 (2016).

74. Wild, J. M. The ventromedial hypothalamic nucleus in the zebra finch (Taeniopygia guttata): Afferent and efferent projections in relation to the control of reproductive behavior. J. Comp. Neurol. 525, 2657–2676 (2017).

75. Harris, K. D. & Shepherd, G. M. G. The neocortical circuit: Themes and variations. Nature Neuroscience vol. 18 170–181 (2015).

76. Braun, K., Scheich, H., Heizmann, C. W. & Hunziker, W. Parvalbumin and calbindin-D28K immunoreactivity as developmental markers of auditory and vocal motor nuclei of the zebra finch. Neuroscience 40, 853–869 (1991).

77. Anderson, S. A., Eisenstat, D. D., Shi, L. & Rubenstein, J. L. R. Interneuron migration from basal forebrain to neocortex: Dependence on Dlx genes. Science (80-.). 278, 474–476 (1997).

78. Kim, T. et al. Cortically projecting basal forebrain parvalbumin neurons regulate cortical gamma band oscillations. Proc. Natl. Acad. Sci. U. S. A. 112, 3535–3540 (2015).

79. Kawaguchi, Y. & Kubota, Y. Correlation of physiological subgroupings of nonpyramidal cells with parvalbumin- and calbindin(D28k)-immunoreactive neurons in layer V of rat frontal cortex. J. Neurophysiol. 70, 387–396 (1993).

80. Phillips, E. A. & Hasenstaub, A. R. Asymmetric effects of activating and inactivating cortical interneurons. Elife 5, (2016).

81. Wood, K. C., Blackwell, J. M. & Geffen, M. N. Cortical inhibitory interneurons control sensory processing. Current Opinion in Neurobiology vol. 46 200–207 (2017).

82. Mahrach, A., Chen, G., Li, N., van Vreeswijk, C. & Hansel, D. Mechanisms underlying the response of mouse cortical networks to optogenetic manipulation. Elife 9, (2020).

83. Medina, L. & Reiner, A. Do birds possess homologues of mammalian primary visual, somatosensory and motor cortices? Trends in Neurosciences vol. 23 1–12 (2000).

84. Saldanha, C. et al. Distribution and Regulation of Telencephalic Aromatase Expression in the Zebra Finch Reveales With a Specific Antibody. J. Comp. Neurol. 423, 619–630 (2000).

85. Ye, J. H., Zhang, J., Xiao, C. & Kong, J. Q. Patch-clamp studies in the CNS illustrate a simple new method for obtaining viable neurons in rat brain slices: Glycerol replacement of NaCl protects CNS neurons. J. Neurosci. Methods 158, 251–259 (2006).

86. Dagostin, A. A., Lovell, P. V., Hilscher, M. M., Mello, C. V. & Leão, R. M. Control of Phasic Firing by a Background Leak Current in Avian Forebrain Auditory Neurons. Front. Cell. Neurosci. 9, (2015).

87. Remage-Healey, L. & Joshi, N. R. Changing Neuroestrogens Within the Auditory Forebrain Rapidly Transform Stimulus Selectivity in a Downstream Sensorimotor Nucleus. J. Neurosci. 32, 8231–8241 (2012).

88. Coleman, M. J. & Mooney, R. Synaptic Transformations Underlying Highly Selective Auditory Representations of Learned Birdsong. J. Neurosci. 24, 7251–7265 (2004).

89. Caras, M., Sen, K., Rubel, E. W. & Brenowitz, E. A. Seasonal Plasticity of Precise Spike Timing in the Avian Auditory System. J. Neurosci. 35, 3431–3445 (2015).

90. Schreiber, S., Fellous, J. M., Whitmer, D., Tiesinga, P. & Sejnowski, T. J. A new correlation-based measure of spike timing reliability. Neurocomputing 52–54, 925–931 (2003).

91. Cohen, J. Statistical Power Analysis for the Behavioral Sciences. (Routledge Academic, 1988).

92. Siegle, J. H. et al. Open Ephys: an open-source, plugin-based platform for multichannel electrophysiology Related content. J. Neural Eng. 14, 045003 (2017).

93. Pachitariu, M., Steinmetz, N., Kadir, S., Carandini, M. & Harris, K. Fast and accurate spike sorting of high-channel count probes with KiloSort.

94. Garcia, S. et al. Neo: an object model for handling electrophysiology data in multiple formats. Front. Neuroinform. 8, (2014).

