## Supplemental figures S1-S7 for "Genetically-identified cell types in avian pallium mirror core principles of excitatory and inhibitory neurons in mammalian cortex"

fig. S1. 60x exemplar images of CaMKIIα (top row; green) and GAD1 (bottom row; magenta) cell morphologies encountered in zebra finch caudomedial nidopallium (NCM), a region of the auditory association pallium. Scale bars all represent 50 microns.

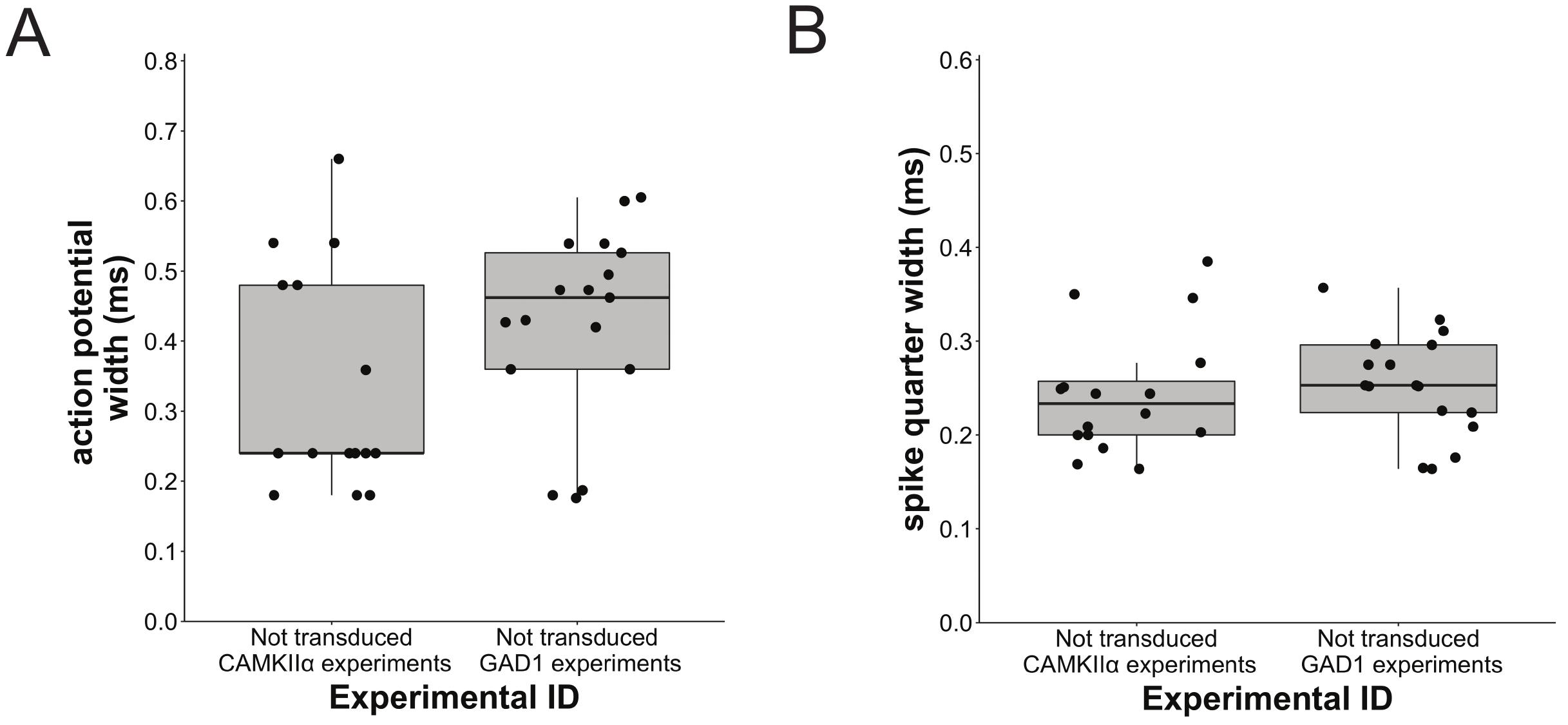

fig. S2. (**A**) Action potential widths and (**B**) spike quarter widths for single units unaffected by light pulses (i.e., not transduced with virus) in CaMKIIα-ChR2 and GAD1-ChR2 optrode experiments *in vivo*. Single units shown here were isolated from the same recording sites as optically-identified single units in Fig. 2G-J.

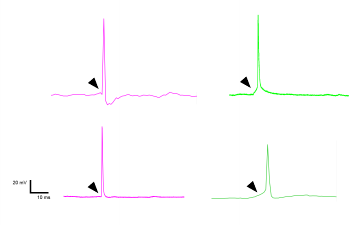

fig. S3. Voltage traces in whole-cell current clamp mode of light-evoked spikes from two representative GAD1-ChR2 cells (magenta; left) and two CaMKIIα-ChR2 cells (green; right). Black arrowheads point to slope of spike onset phase.

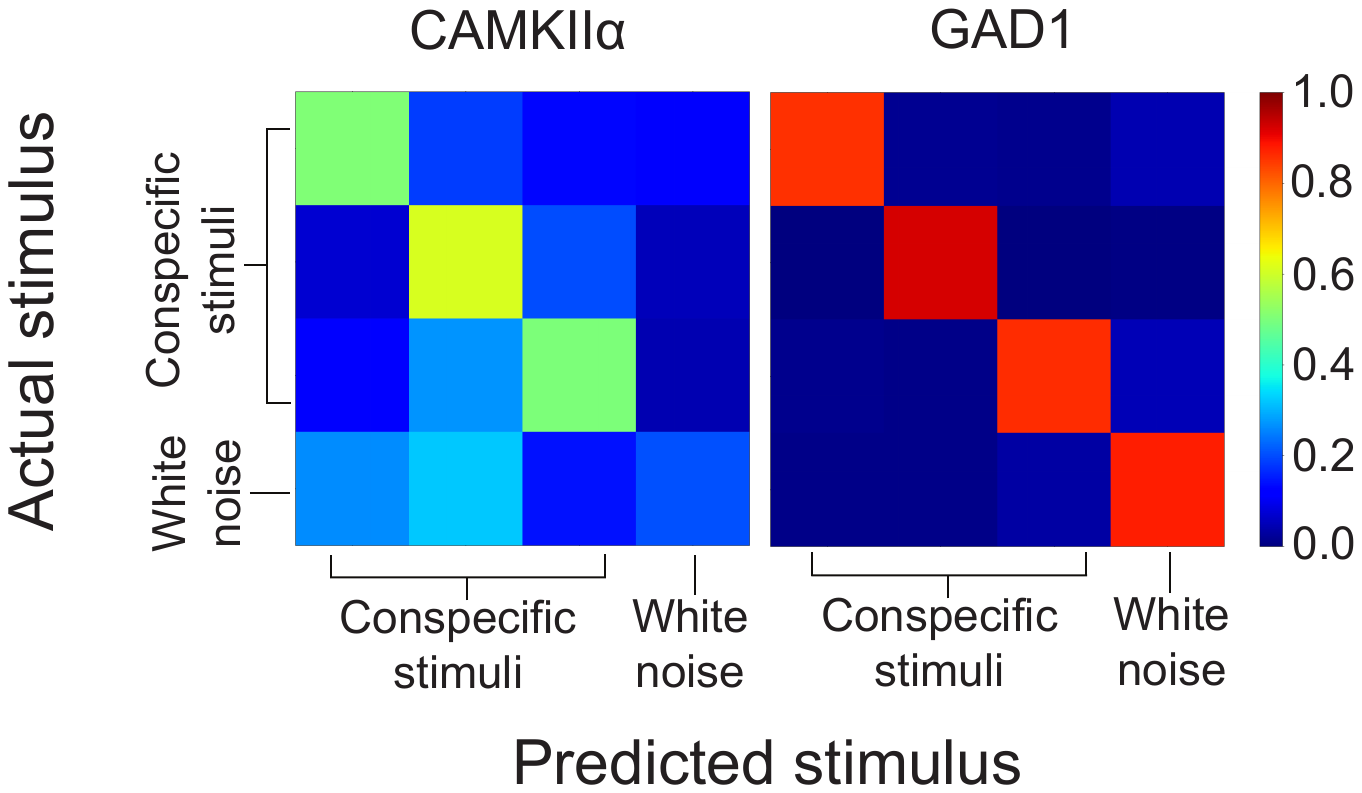

fig. S4. Pattern classifier timing accuracy confusion matrices for a representative CaMKIIα single unit and a GAD1 single unit. The diagonals of the matrices show the degree of accuracy values for the correct associated auditory stimulus predicted by the pattern classifier from single unit spike trains. Values closer to 1 on the heat map represent higher timing accuracy.

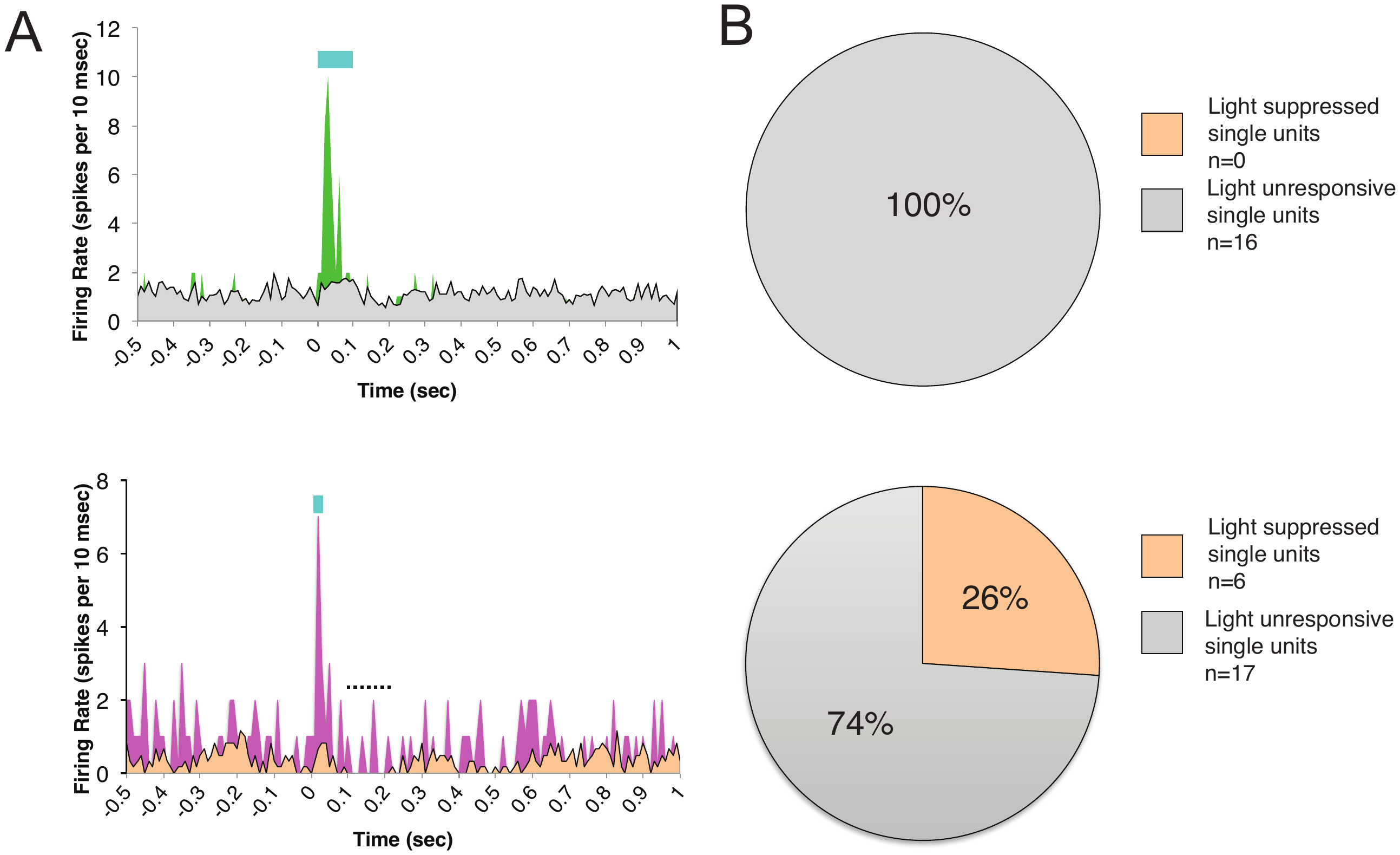

fig. S5. Evidence of feedforward suppression in GAD1-ChR2 optrode recordings but not CaMKIIα-ChR2 optrode recordings. (**A**) Firing rates of an exemplar CaMKIIα-ChR2 transduced single unit (top) and GAD1-ChR2-transduced single unit (bottom) in response to light pulse. Top: Gray trace represents average firing rate for non-transduced single units in CaMKIIα-ChR2 optrode recordings, which showed no response to light. Bottom: Orange trace represents average firing rate for a subset of non-transduced single units in GAD1-ChR2 optrode recordings, which demonstrated suppression following light pulses. (**B**) Pie charts demonstrating fraction of light-suppressed non-transduced units in CaMKIIα-ChR2 (top) and GAD1-ChR2 (bottom) optrode recordings.

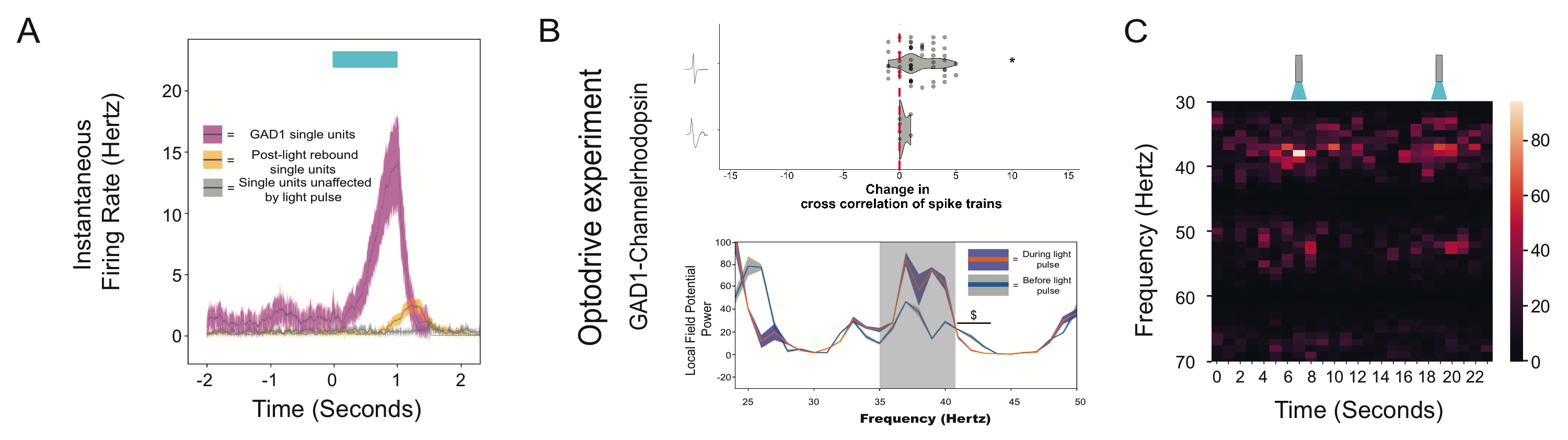

fig. S6. *In vivo* GAD1-ChR2 optodrive experiments in in an awake male zebra finch (N = 1). (**A**) Instantaneous firing rates of single units that increased firing rate in response to 1 sec light pulse (magenta; n = 3), single units that increase in firing rate after 1 sec light pulse (orange; n = 9), and light unresponsive single units (gray; n = 7) drawn from n = 30 total units isolated in NCM. (**B**) (Top) Violin plot shows change in cross-correlation of waveforms following stimulation with blue light. Red line denotes zero change. (Bottom) Local field potential (LFP) power spectra before and during stimulation of NCM with blue light. Grey shading represents gamma frequency range for which LFP power is significantly higher compared to baseline (P<0.05); $ is gamma frequency range for which LFP power is significantly lower than baseline (P<0.05). (**C**) Heatmap (% max LFP power) of an exemplar change over time in response to a pair of blue light pulses.

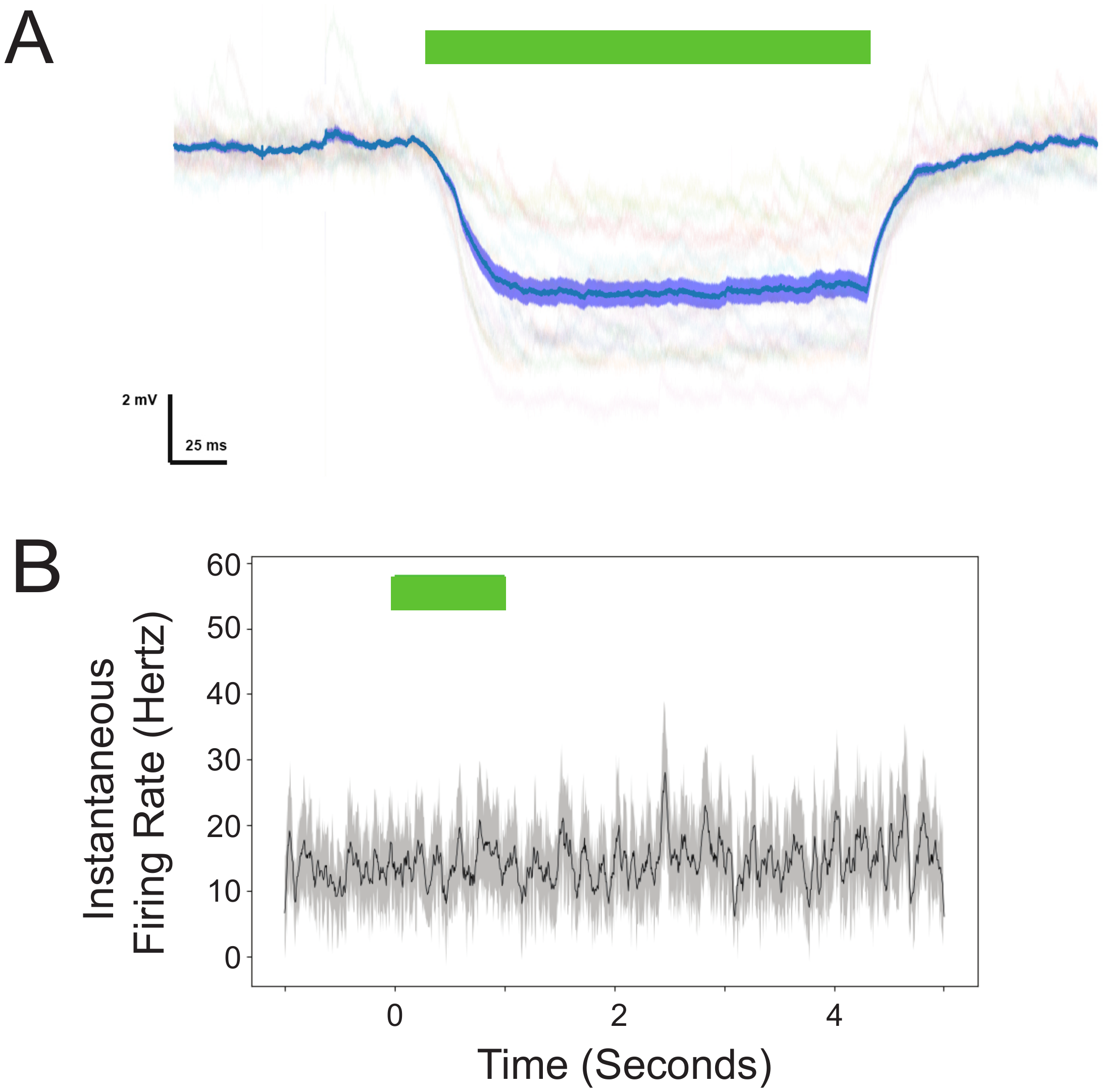

fig. S7. GAD1-archaerhodopsin stimulation elicits hyperpolarizing currents specifically in cells transduced with the virus. (**A**) Voltage trace in whole-cell current clamp of a cell transduced with GAD1-archaerhodopsin. Turquoise trace represents mean membrane voltage (deflections from resting membrane potential) across several trials (individual trials shown as faint background traces) and purple shading is ± 1 SEM. Green line above indicates duration of 525 nm light pulse. (**B**) Exemplar trace of n = 4 non-transduced single units from a site in NCM that also contained transduced units. Green bar shows 1 sec green (525 nm) light pulse.
